# RAID: Regression Analysis based Inductive DNA microarray for Precise Read-Across

**DOI:** 10.1101/2022.02.15.480621

**Authors:** Yuto Amano, Masayuki Yamane, Hiroshi Honda

**Affiliations:** R&D Safety Science Research, Kao Corporation, 2606 Akabane, Ichikai-Machi, Haga-Gun, Tochigi, Japan

**Author notes:** **Correspondence:** Hiroshi Honda, Ph.D.

## Abstract

Chemical structure-based read-across represents a promising method for chemical toxicity evaluation without the need for animal testing; however, a chemical structure is not necessarily related to toxicity. Therefore, *in vitro* studies were often used for read-across reliability refinement; however, their external validity has been hindered by the gap between *in vitro* and *in vivo* conditions. Thus, we developed a virtual DNA microarray, <u>R</u>egression <u>A</u>nalysis based Inductive <u>D</u>NA microarray (RAID), which quantitatively predicts *in vivo* gene expression profiles based on the chemical structure and/or *in vitro* transcriptome data. For each gene, elastic-net models were constructed using chemical descriptors and *in vitro* transcriptome data to predict *in vivo* data from *in vitro* data (*in vitro* to *in vivo* extrapolation; IVIVE). In feature selection, useful genes for assessing the quantitative structure activity relationship (QSAR) and IVIVE were identified. Predicted transcriptome data derived from the RAID system reflected the *in vivo* gene expression profiles of characteristic hepatotoxic substances. Moreover, gene ontology and pathway analyses indicated that xenobiotic response and metabolic activation via nuclear receptors are related to those gene expressions. The identified IVIVE-related genes were associated with fatty acid-, xenobiotic-, and drug metabolism, indicating that *in vitro* studies were effective in evaluating these key events. Furthermore, validation studies revealed that chemical substances associated with these key events could be detected as hepatotoxic biosimilar substances. These results indicate that the RAID system could represent an alternative screening test for repeated-dose toxicity test and toxicogenomic analyses. Our technology provides a critical solution to IVIVE-based read-across by considering the mode of action and chemical structures.

## 1 Introduction

Non-animal testing for efficacy and safety evaluation of chemical substances is one of the key concepts of balancing animal welfare and efficient development. Since the marketing ban in the EU in March 2013 ((EC) No. 1223/2009) (EU, 2009) of cosmetic products and ingredients tested on animal models, safety assessment methodologies independent of animal testing have attracted much attention. Simultaneously, the utilization of non-animal high-throughput technology for optimizing drug discovery processes is becoming highly important in pharmaceuticals (Loiodice et al., 2017; Rognan, 2017; Amano et al., 2020).

Read-across, a process that estimates substance toxicity based on the concept that substances with similar chemical structure have similar biological activity, represents a promising approach and has already been conceptually accepted as a reliable safety risk assessment by some regulatory authorities (ECHA, 2017; European Commission, 2018). Likewise, quantitative structure activity relationship (QSAR) has been widely used and impurity characterization received regulatory acceptance (ICH M7). However, since subtle structural differences may elicit different biological responses, supporting the read-across robustness by using biological similarities has been considered important (Ball et al., 2016, 2020; Zhu et al., 2016). Registration, Evaluation, Authorization, and Restriction of Chemicals (REACH) mentioned that the read-across performed by registrants often fail to comply with the legal requirements due to defects in the hypothesis and justification of the toxicological prediction (ECHA, 2020).

There are two approaches to enhance the reliability of read-across: (1) Employment of *in vitro* data relevant to specific toxicity. Methodologies to incorporate *in vitro* data within read-across (Ball et al., 2016, 2020; ECHA, 2017; Guo et al., 2019) and some case studies (OECD, 2016a, 2016b, 2018; Nakagawa et al., 2020, 2021) have been reported. However, these approaches can be applied only to specific toxicity endpoint and substances with a known toxicity and mode of action. Such conditions were previously termed as “local validity” (Patlewicz et al., 2014). (2) The use of biologically similar substances based on their profiles obtained from a large number of bioassays. The United States Environmental Protection Agency’s (US EPA’s) research project, ToxCast and Tox21, provided hundreds of high-throughput screening assays and several groups employed such biological activity data for toxicological evaluation (Sipes et al., 2013; Berggren et al., 2015; Richard et al., 2021). Although this concept could be applied to substances with little information to elucidate their entire toxicological profiles and find their key mode of action, it is time-consuming and expensive to conduct numerous bioassays for a new candidate substance. In contrast, transcriptome data containing approximately 30,000 gene expression values can be used to estimate perturbated mechanisms through enrichment analysis. Wang et al. (2016) tried to predict drug-induced adverse effects by employing LINCS L1000 data (Subramanian et al., 2017), whereas Iwata et al. (2019) developed a computational method to predict missing value from the LINCS L1000 transcriptomic profiles of various human cell lines and provided new drug therapeutic indications. Genomic data have been considered to be usable in read-across by Health Canada and a research group from the U.S. FDA (Health Canada, 2019; Liu et al., 2019). However, several researchers showed that *in vitro* gene expression values are not always highly correlated with *in vivo* data (Sutherland et al., 2016; Grinberg et al., 2018; Liu et al., 2018). Thus, interpreting toxicological meaning from the *in vitro*-*in vivo* relationship and *in vitro* to *in vivo* extrapolation (IVIVE) in omics data represents a big challenge for chemical risk assessment.

As an IVIVE study in omics data, Liu et al. (2020) developed a useful *in silico* strategy to narrow the data gap between *in vitro* and *in vivo* conditions. They modified *in vitro* data using non-generative matrix factorization methods to improve the correlation with *in vivo* data, which overcame the shortcomings of previous large-scale genomic data predictions regarding the *in vitro*-*in vivo* data gap (Liu et al., 2020). Although non-generative matrix factorization enables macroscopic estimation based on a pattern recognition classifying chemical and biological responses, it does not focus on each gene estimation. As an alternative solution, microscopic estimation for each gene expression were performed based on tensor-train weighted optimization using machine learning (Iwata et al., 2019); however, such comprehensive estimation have not been integrated within an IVIVE study. Therefore, predicting *in vivo* transcriptomic profiles from *in vitro* data for IVIVE might not only enhance the robustness of read-across but could also be utilized in other non-animal testing strategies as weight of evidence, such as in Integrated Approaches to Testing and Assessment (IATA) and New Approach methods (NAMs) for safety and drug repositioning research.

In this study, we developed a virtual DNA microarray that quantitatively predicts the *in vivo* gene expression profiles based on the chemical structure and/or *in vitro* transcriptome data. For each gene, elastic-net models, a regression analysis method that has been used in toxicity prediction with visualization of feature importance (e.g. Fujita et al., 2020), were constructed using chemical descriptors and *in vitro* transcriptome data. We named the set of prediction models “Regression Analysis based Inductive DNA microarray (RAID)” to inductively analyze the mode of action and the key event in adverse effects with reference to the Redundant Arrays of Inexpensive Disks, a data storage virtualization technology also represented as RAID that combines multiple physical disk drive components with the purpose of data redundancy. As RAID (storage technology) complements data based on the information of multiple components, we hope that RAID (our microarray) will complement the relationships between multiple media (*in vivo* gene expression, *in vitro* gene expression, and chemical structure). RAID system achieved the quantitative *in vitro* to *in vivo* extrapolation (QIVIVE) by the integration of a structure-based approach (QSAR) with transcriptomic data. Whereas general “Q”IVIVE studies predict dose (or concentration) quantitatively in toxicological or toxicokinetic effects, our “Q”IVIVE predicts *in vivo* gene expression values quantitatively. Finally, the substance similarities were analyzed by principal component analysis (PCA), which proved useful in understanding the features of toxic substances based on their gene expression profile (Watanabe et al., 2012), using RAID (the virtual microarray) data, *in vivo* data, *in vitro* data, and chemical structure data to validate the usefulness of read-across.

## 2 Materials and Methods

### 2.1 Gene expression and chemical structure data

No animal experiment has been performed in this study. The transcriptome data from DNA microarrays (Affymetrix Rat Genome 230 2.0 chips; Santa Clara, CA, USA) were extracted from the Toxicogenomics Project-Genomics Assisted Toxicity Evaluation system (TG-GATEs). TG-GATEs contains *in vitro* and *in vivo* transcriptome data for rat single- and repeated-dose toxicity tests of 170 compounds (Igarashi et al., 2015). The transcriptome data obtained from the livers of rats treated with high doses for 28 days and primary rat hepatocytes treated with high doses for 24 h were downloaded and pre-processed using MAS5 (Gautier et al., 2004). In this study, chemical substances tested *in vitro* and *in vivo*, that fulfilled a maximum sample number (n = 2 for *in vitro* and n = 3 for *in vivo*), and had no incalculable chemical descriptors (described below), were analyzed. Thus, 115 compounds were examined in this study (Table 1).

**Table 1.** List of compounds used in the present study and their toxicological classes.

| Tox<br>class<sup>1)</sup> | Compound name |
| --- | --- |
| Toxic | Allyl alcohol (AA), 2-acetamidofluorene (AAF), α-naphthyl isothiocyanate (ANIT), Acetaminophen (APAP), Aspirin (ASA), Benzbromarone (BBr), Bromobenzene (BBZ), Bucetin (BCT), Bendazac (BDZ), Benziodarone (BZD), carboplatin (CBP), Coumarin (CMA), Chlormezanone (CMN), Chloramphenicol (CMP), Colchicine (COL), Cyclophosphamide monohydrate (CPA), Clomipramine hydrochloride (CPM), Chlorpropamide (CPP), Cyclosporine A (CPA), Diltiazem hydrochloride (DIL), Disopyramide (DIS), Disulfiram (DSF), Dantrolene sodium hemiheptahydrate (DTL), Diazepam (DZP), Ethambutol dihydrochloride (EBU), 17-α-ethinylestradiol (EE), DL-ethionine (ET), Fenofibrate (FFB), Flutamide (FT), Gemfibrozil (GFZ), Hexachlorobenzene HCB), Lomustine (LS), Mexiletine hydrochloride (MEX), Methapyrilene hydrochloride (MP), Methyltestosterone (MTS), Methimazole (MTZ), Nimesulide (NIM), Phenacetin (PCT), Promethazine hydrochloride (PMZ), Propylthiouracil (PTU), Sulfasalazine (SS), Simvastatin (SST), Sulindac (SUL), Thioacetamide (TAA), Terbinafine hydrochloride (TBF), Ticlopidine hydrochloride (TCP), Trimethadione (TMD), Vitamin A (VA), WY-14643 (WY) |
| Non-toxic | Acarbose (ACA), Acetazolamide (ACZ), Adapin (ADP), Ajmaline (AJM), Amiodarone hydrochloride (AM), Amitriptyline hydrochloride (AMT), Allopurinol (APL), 2-bromoethylamine hydrobromide (BEA), Caffeine (CAF), Captopril (CAP), Carbamazepine (CBZ), Clofibrate (CFB), Chlorpheniramine maleate (CHL), Cimetidine (CIM), Chlormadinone acetate (CLM), Cephalothin sodium (CLT), Ciprofloxacin hydrochloride (CPX), Chlorpromazine hydrochloride (CPZ), Diclofenac sodium (DFNa), Danazol (DNZ), Erythromycin ethylsuccinate (EME), Enalapril maleate (ENA), Ethanol (ETN), Etoposide (ETP), Famotidine (FAM), Fluphenazine dihydrochloride (FP), Furosemide (FUR), Glibenclamide (GBC), Griseofulvin (GF), Gentamicin sulfate (GMC), Haloperidol (HPL), Hydroxyzine dihydrochloride (HYZ), Ibuprofen (IBU), Imipramine hydrochloride (IMI), Isoniazid (INAH), Iproniazid phosphate (IPA), Ketoconazole (KC), Methyldopa (MDP), Mefenamic acid (MEF), Metformin hydrochloride (MFM), Moxisylyte hydrochloride (MXS), Nitrofurantoin (NFT), Nitrofurazone (NFZ), Nicotinic acid (NIC), Nifedipine (NIF), Omeprazole (OPZ), Papaverine hydrochloride (PAP), Phenobarbital sodium (PB), D-penicillamine (PEN), Perhexiline maleate (PH), Phenylbutazone (PhB), Phenytoin (PHE), Pemoline (PML), Quinidine sulfate (QND), Ranitidine hydrochloride (RAN), Rifampicin (RIF), Sulpiride (SLP), Tannic acid (TAN), Tetracycline hydrochloride (TC), Tiopronin (TIO), Tolbutamide (TLB), Tamoxifen citrate (TMX), Triamterene (TRI), Thioridazine hydrochloride (TRZ), Triazolam (TXM), Sodium valproate (VPA) |

1) The toxicological classes of chemical substances were referred to a previous report (Low et al., 2011). The authors classified these compounds into histopathological and serum chemistry classes. Compounds with hepatotoxic histopathological findings and other histopathological findings with biochemical marker changes in serum chemistry were defined toxic-compounds in this study.

For the chemical structure data, the alvaDesc chemical descriptors (Mauri, 2020) were calculated using alvaDesc v1.0 software (Alvascience-Srl, Lecco, Italy). AlvaDesc can calculate 3885 2D-descriptors and 1420 3D-descriptors. However, only 2D-descriptors were used excluding those with a high pair correlation (>0.95), constant for all substances, and at least one missing value. Consequently, 854 descriptors were calculated. Each descriptor was normalized using the bestNormalize package (ver. 1.8.0) in R (ver. 4.1.1) (https://cran.r-project.org/). This package estimates the optimal normalizing transformation from Yeo-Johnson transformation, the Box Cox transformation, the log_10_ transformation, the square-root transformation, and the arcsine transformation.

### 2.2 Construction of the RAID system (a virtual microarray)

To extrapolate *in vitro* transcriptome data to *in vivo* conditions, we developed predictive models for each gene. The predictive models predicting *in vivo* transcriptome data from chemical descriptors and *in vitro* data were developed using the elastic-net regression method. The value of each cell in the matrix was the fold change on a base 2 logarithmic scale. The set of those predictive models was named a virtual microarray “RAID” (as mentioned in the Introduction) (Figure 1). To suppress over-learning, the hyperparameters (α and λ) of each model were optimized with a 5-fold cross-validation. We removed the genes that were associated with less than 10 chemical substances inducing differential expression (<1.5 fold change) since it would be difficult to run machine learning scripts on such rare genes. Consequently, RAID was composed of 1601 prediction models for each gene.

**Figure 1.**
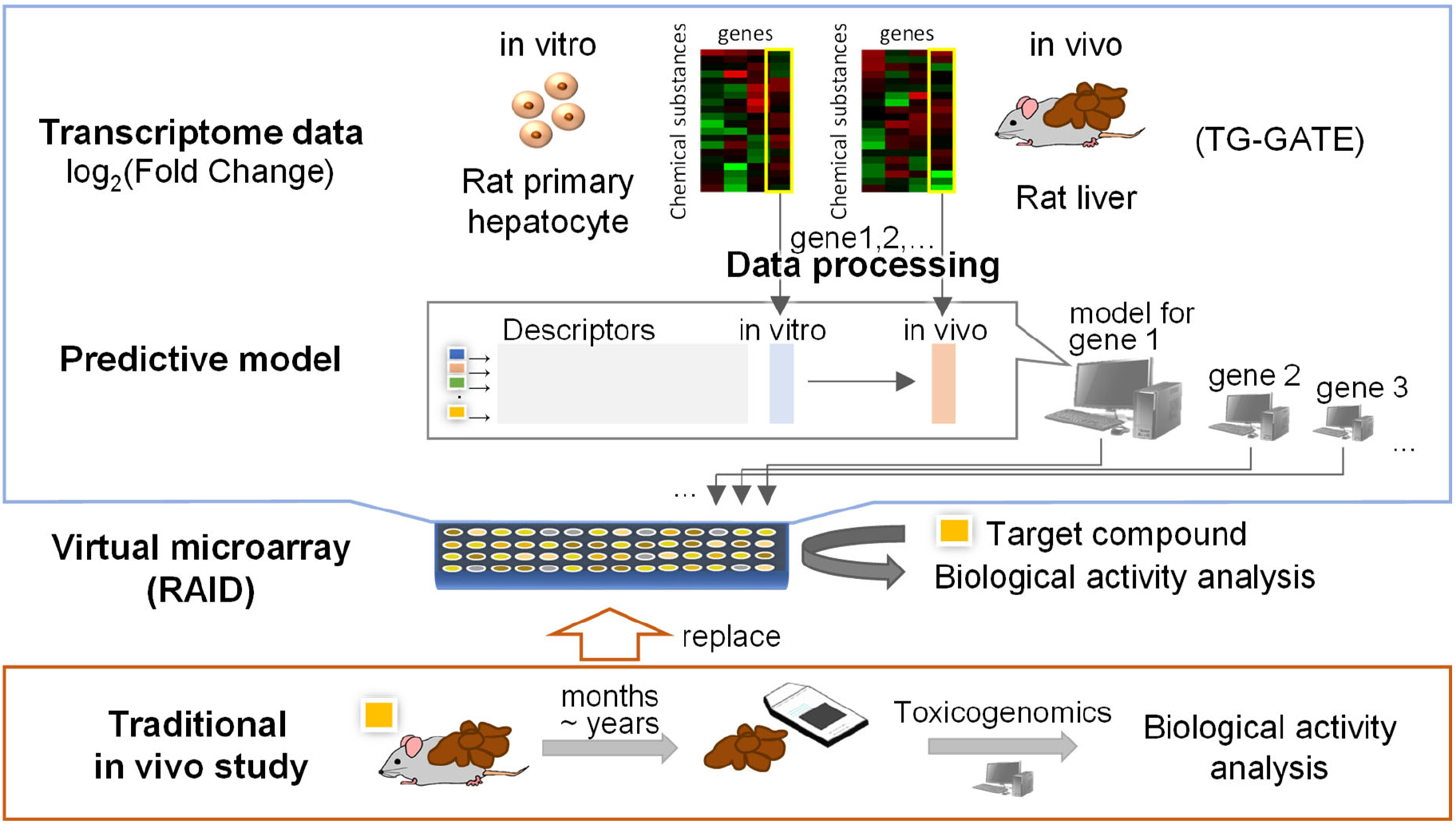
Approach to construct a virtual microarray (RAID). The predictive model for comprehensive *in vivo* transcriptome data was constructed using elastic-net regression as well as chemical descriptors and *in vitro* transcriptome data.

To construct RAID that correctly predicts the bioactivities of chemical substances, the quality of training data sets was extremely important, and differentially expressed genes should be determined strictly considering data noise. Hence, we addressed this issue by data processing (feature engineering) and model justification. First, after calculating the fold change values (sample treated groups/solvent control group), the gene differentiation values with low reliability were adjusted. Briefly, the fold change value increments were changed to half (e.g. 1.5 decreased to 1.25) in the sample with the number of flag A (low reliability) ≥ 2 out of 3 for *in vivo* and the number of flag A ≥ 1 out of 2 for *in vitro*, or in the sample with p-value ranging between 0.05 and 0.1. The fold change values were changed one-fourth (e.g. 1.4 decreased to 1.1) in the sample with p-value over 0.1, and were treated as 1 (no differentiation) in the sample with flags all A in both *in vivo* and *in vitro*. Second, weight parameters were used in model building. The weight of samples with ≥1.5 fold change was set to 1.5 and ≥4 fold change was set to 2.

### 2.3 Interpretation of biological meaning of RAID analysis

Considering the application of RAID to read-across, the gene expression data was visualized by PCA using prcomp function from stats package (ver. 4.1.1) and probability ellipse frames of toxic and non-toxic substances were drawn using the ggfortify package (ver. 0.4.12) in R to compare *in vivo*, *in vitro*, and chemical descriptor data. The toxic class of chemical substances were determined based on previously reported histopathological and serum chemistry findings (Table 1) (Low et al., 2011). As a reference data point, the biological meaning of genes that contributed to the PCA plot of *in vivo* data was analyzed using pathway analysis. The loading value of genes in the PCA was defined as length of loadings calculated using Pythagorean theorem

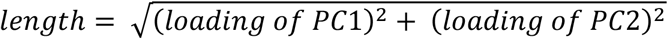

and genes with top 30 loading value in 1^st^ and 4^th^ quadrant were analyzed.

To analyze the biological consistency with *in vivo* data, commonality of principal component related genes (top and bottom 30 rotations in each PC1 and PC2 of PCA) were visualized using the VennDiagram package (ver. 1.6.20) in R, and enrichment analyses of each categorized gene were conducted using Gene Ontology-biological process and Reactome pathway by Metascape (Zhou et al., 2019). Four categorized genes related to *in vivo* data (*in vivo* only, *in vivo* and RAID, *in vivo* and *in vitro*, and all three data) were analyzed to characterize which biological process could be covered by RAID and *in vitro* data. Furthermore, to characterize genes whose predictive models in RAID used *in vitro* data, enrichment analysis of top 20 genes with the highest importance (contribution) for *in vitro* data in the model was conducted. In the analysis, Affymetrix probe ID was converted to gene symbol using the biomaRt package (ver. 2.50.2) in R.

### 2.4 Quantitative IVIVE effects in RAID system

For performance evaluation against the quantitative IVIVE, root-mean-square errors (RMSEs) of RAID predicted values to *in vivo* data were calculated and compared to those of *in vitro* data. To exclude the difference in gene expression value distribution of each data source, fold change values were normalized before RMSEs was calculated. The RMSEs were calculated for both all genes and genes for which *in vitro* data had importance in the model.

### 2.5 Read-across application using external data

To validate the usefulness of RAID for functional read-across-based analysis of both predicted gene expression profiles and chemical structures, substances that did not contain training data sets for model building (Table 1) were further explored using Ingenuity Pathway Analysis (IPA) (QIAGEN Inc., https://www.qiagenbioinformatics.com/products/ingenuitypathway-analysis). Specifically, substances that may promote the expression of genes (have known relationship with the genes) that were identified by the PCA and pathway analysis of *in vivo* data (see section **2.3**) were explored using IPA. Chemical descriptors of each substance were analyzed using alvaDesc v1.0 software (Alvascience-Srl, Lecco, Italy) and gene expression profiles were fulfilled using median values of training data sets. Finally, RAID analyses using constructed predictive models for those substances and re-analyzed PCA data were used to evaluate similarities based on predicted-biological responses.

## 3 Results

### 3.1 Biological analysis of RAID compared to that of *in vivo* and *in vitro* microarray data

RAID (predicted transcriptome) data was visualized using PCA (Figure 2). From a higher perspective, two directions mainly composed of toxic substances were identified and many toxic substances were separated from non-toxic substances via RAID and *in vivo* data, whereas they could not be separated based on *in vitro* and chemical descriptor data. Moreover, two common toxic-substances groups (e.g. 1^st^ group [TAA, MP, and HCB] placed in 1^st^ quadrant and 2^nd^ group [WY, FFB, BBr, and GFZ] placed in 2^nd^ quadrant) were distanced from non-toxic substances along PC1 and PC2 in both RAID and *in vivo* data, nonetheless the PC1 and PC2 replaced. The loading plot showed that *Cyp1a1 (Cytochrome P450, family 1, subfamily A, polypeptide 1)*, *Gpx2 (Glutathione peroxidase 2)*, and *Gsta3 (Glutathione S-transferase A3)* gene expression were commonly observed in RAID and *in vivo* data, and enabled the discrimination of TAA, MP, and HCB. Furthermore, *Acot1 (Acyl-CoA thioesterase 1)*, *Vnn1 (Vanin1),* and *Cyp4a11 (Cytochrome P450, family 4, subfamily A, polypeptide 11)* contributed to discriminating WY, FFB, BBr, and GFZ.

**Figure 2.**
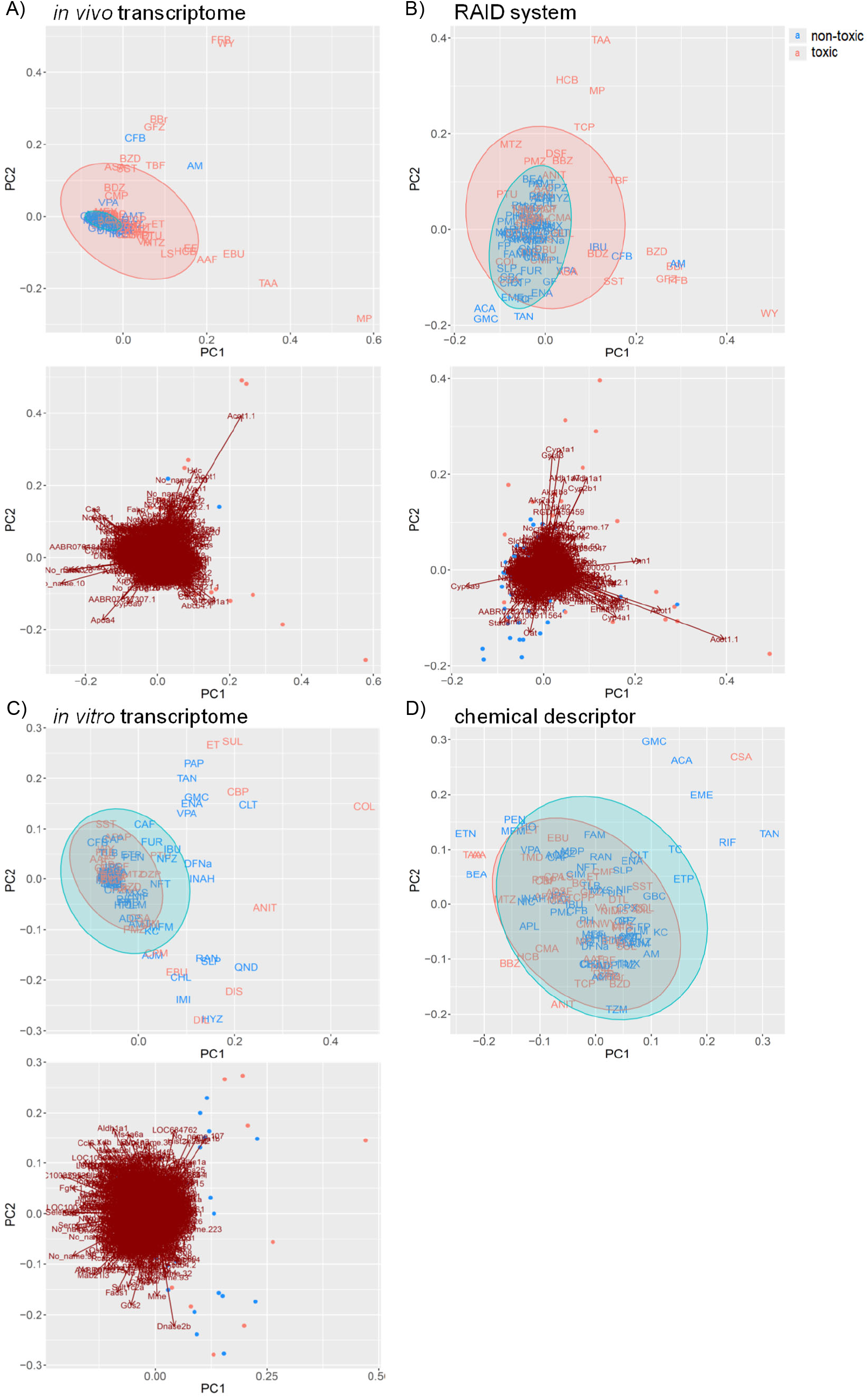
PCA score plots for chemical substances and the gene loading in the transcriptome data of A) *in vivo*, B) virtual microarray (RAID), and C) *in vitro* data. PCA score plot with D) chemical descriptor data. Uppercase letters in PCA score plots: abbreviations of chemical substances are described in Tab.1. Color 1: non-toxic substances. Color 2: hepatotoxic substances. Gene symbols are presented on the arrowhead (loading).

Pathway analysis indicated that the 1^st^ group related genes would be associated with peroxisome proliferative activity characterized by *Cyp4a* induction via peroxisome proliferator-activated receptor-alpha (PPARa) activation and 2^nd^ group related genes would be associated with xenobiotic response, including *Cyp1a* induction via aryl hydrocarbon receptor (AHR) and carcinogenesis (Figure 3). To clarify the biological functions that RAID covers, the commonalities between related genes and principal components were explored (Figure 4A and Table 2). As expected from Figure 2, RAID shared more genes (36; Table 2) with the *in vivo* data than with the *in vitro* data (9). Enrichment analysis revealed that the biological processes related to metabolism and detoxification and pathways associated with peroxisomal protein transport were enriched in both *in vivo* and RAID data, indicating that RAID could cover these functions, and ultimately indicate key functions through pathway analysis (Figure 3). Conversely, although several metabolic processes were enriched within the *in vitro* data, those biological functions were covered by RAID as well (Figure 4B). These results suggest that RAID data allow the detection of more *in vivo* key toxic events than *in vitro* transcriptome data.

**Figure 3.**
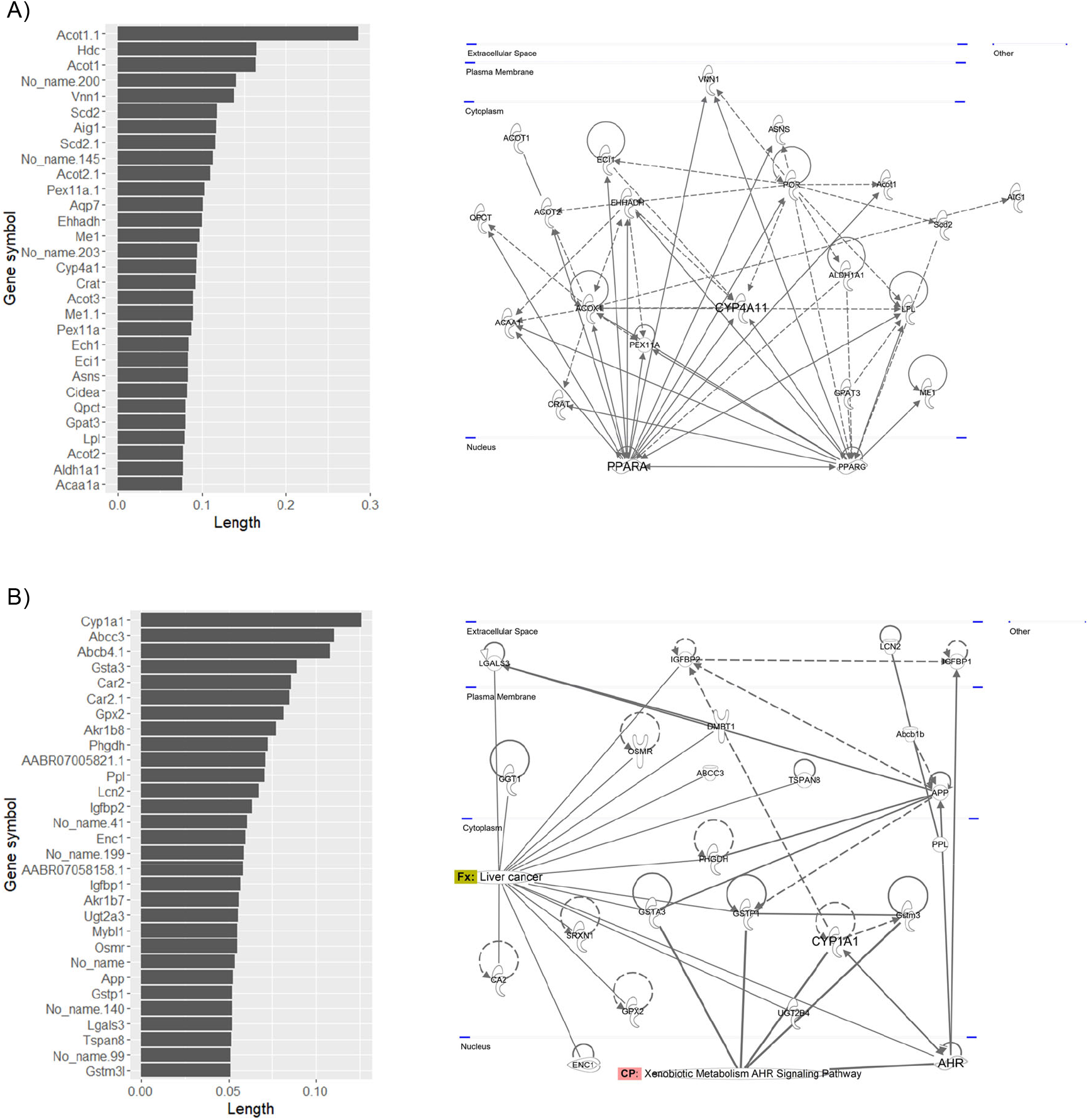
List of genes that have high loading values in the PCA plot of *in vivo* data and their pathway map. The loading value was defined as the loading length in the 1^st^ or 2^nd^ quadrant calculated using the Pythagorean theorem. The pathway map was drawn by upstream regulator analysis using IPA.

**Figure 4.**
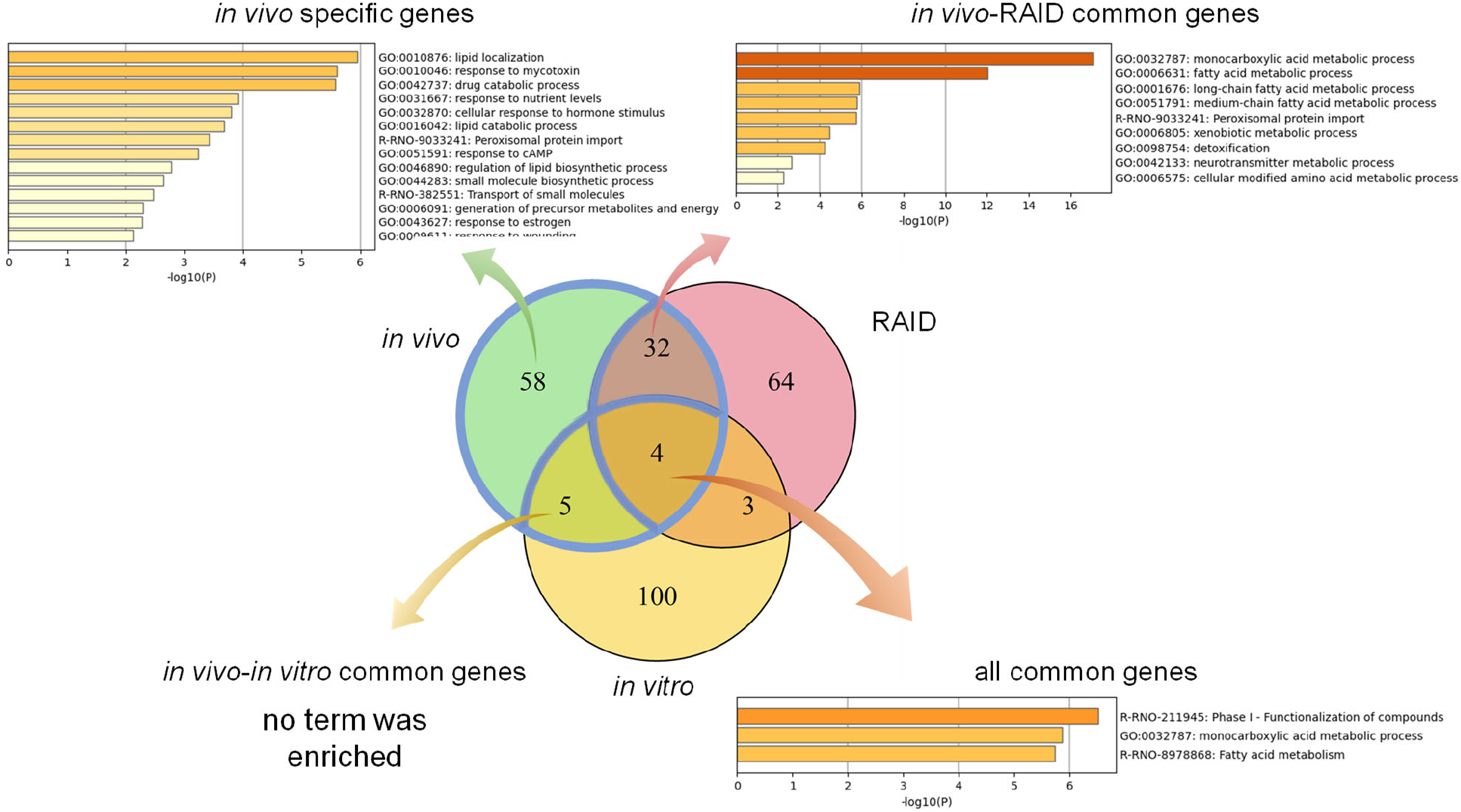
Commonalities of principal components related genes and their biological functions analyzed by gene ontology and pathway analyses. Venn diagram of genes related to the 1^st^ and 2^nd^ principal components of *in vivo*, a virtual microarray (RAID), and *in vitro* data.

**Table 2.**
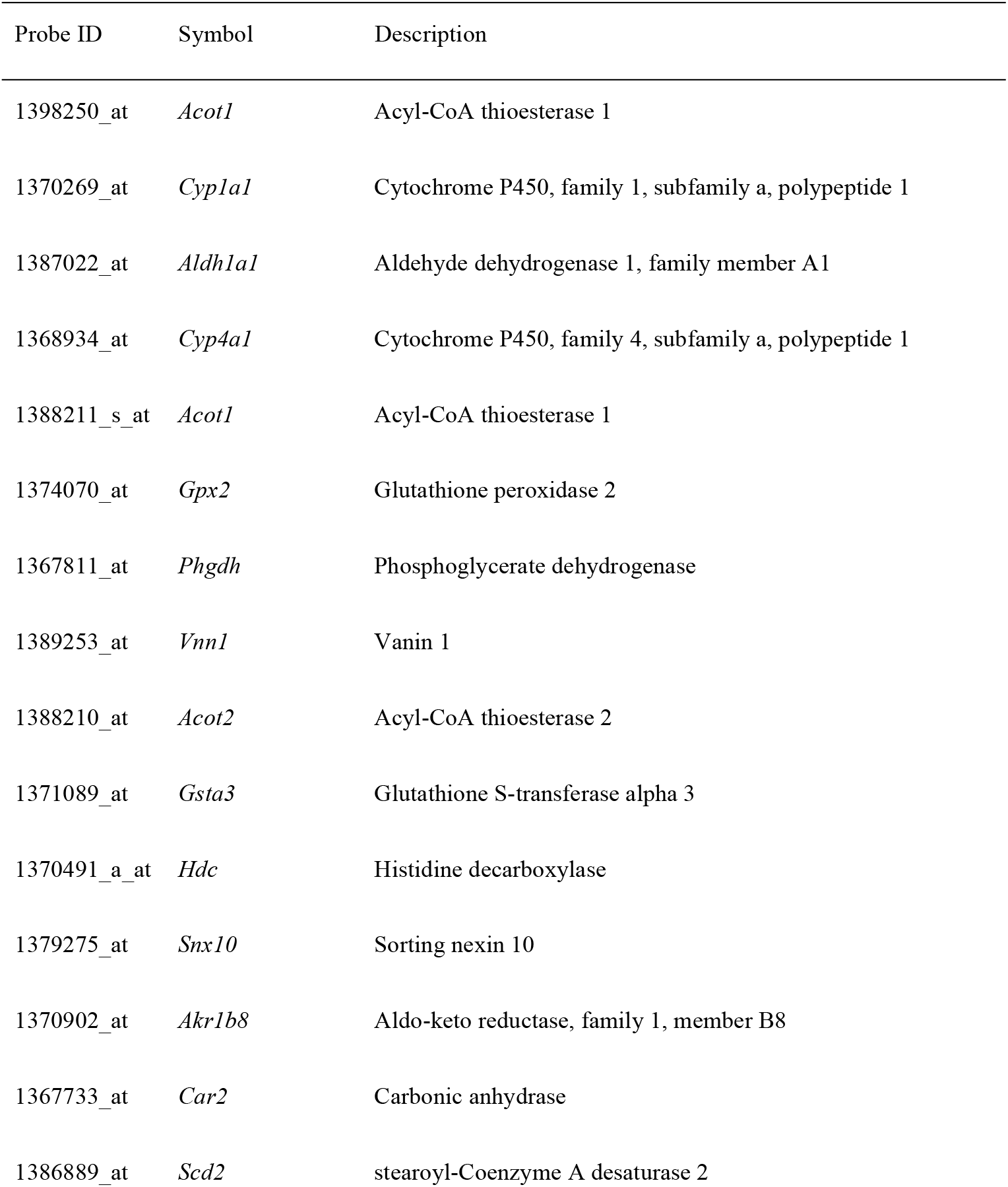
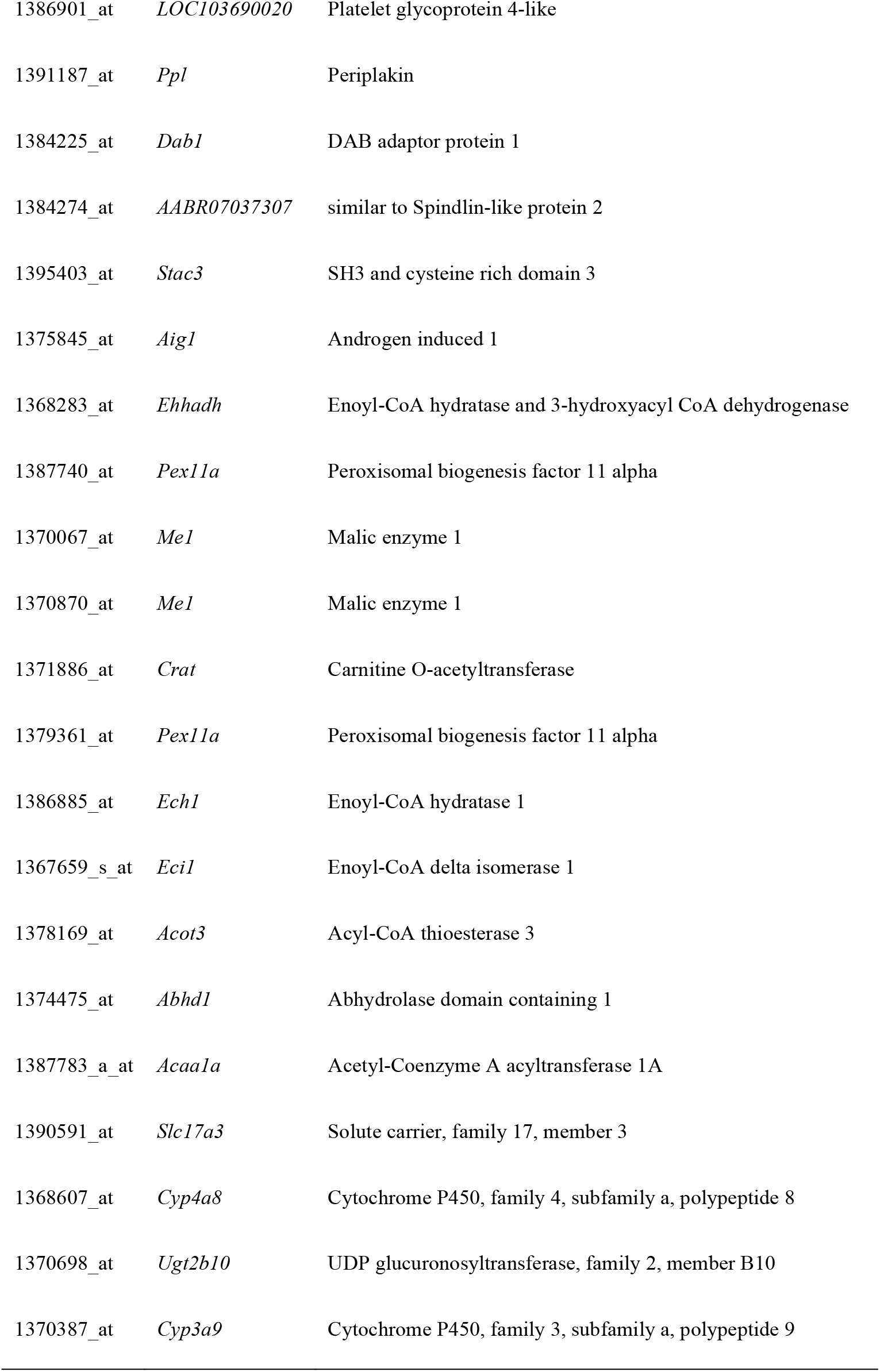
Principal components relating common genes in a virtual microarray (RAID) and in vivo data.

### 3.2 Importance of *in vitro* data in the RAID system

Enrichment analysis of genes whose predictive model used highly relevant *in vitro* data (top 20 genes for which *in vitro* data had high importance in all predictive models; Table 3) indicated that *in vitro* data contributed to estimating the gene expression values associated with metabolic processes of fatty acid, xenobiotics, and drugs, and peroxisome proliferative activity (Pathway on peroxisome protein import and biological process associated with the regulation of peroxisome size; Figure 5).

**Figure 5.**
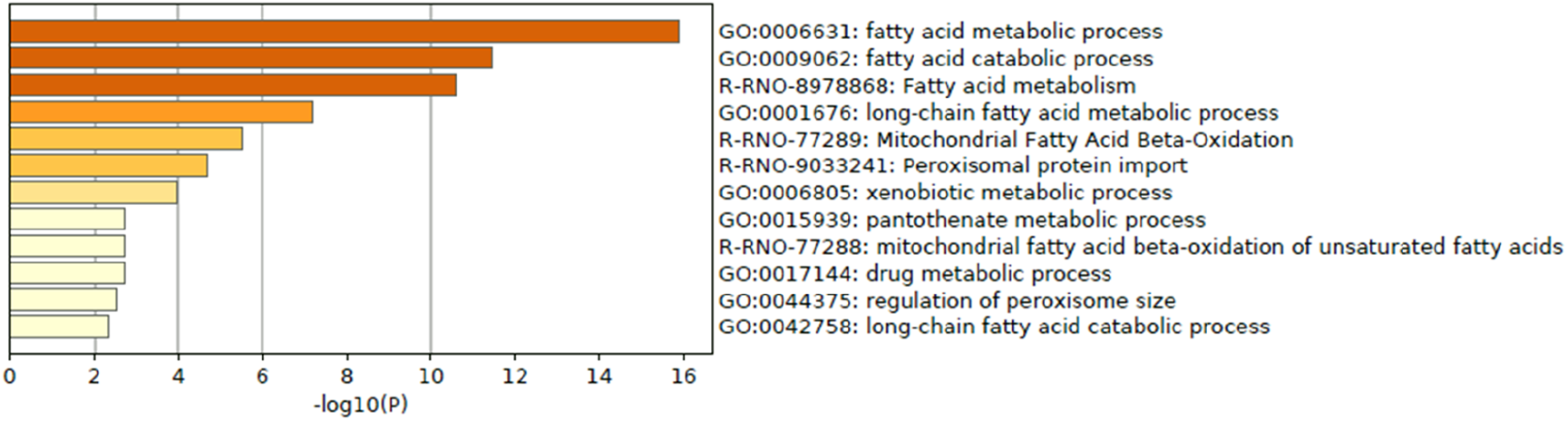
Enrichment analysis of *in vitro-in vivo* extrapolation (IVIVE) related genes identified in a virtual microarray (RAID) system. Top 20 most important (contribution) genes from the predictive models were analyzed.

**Table 3.**
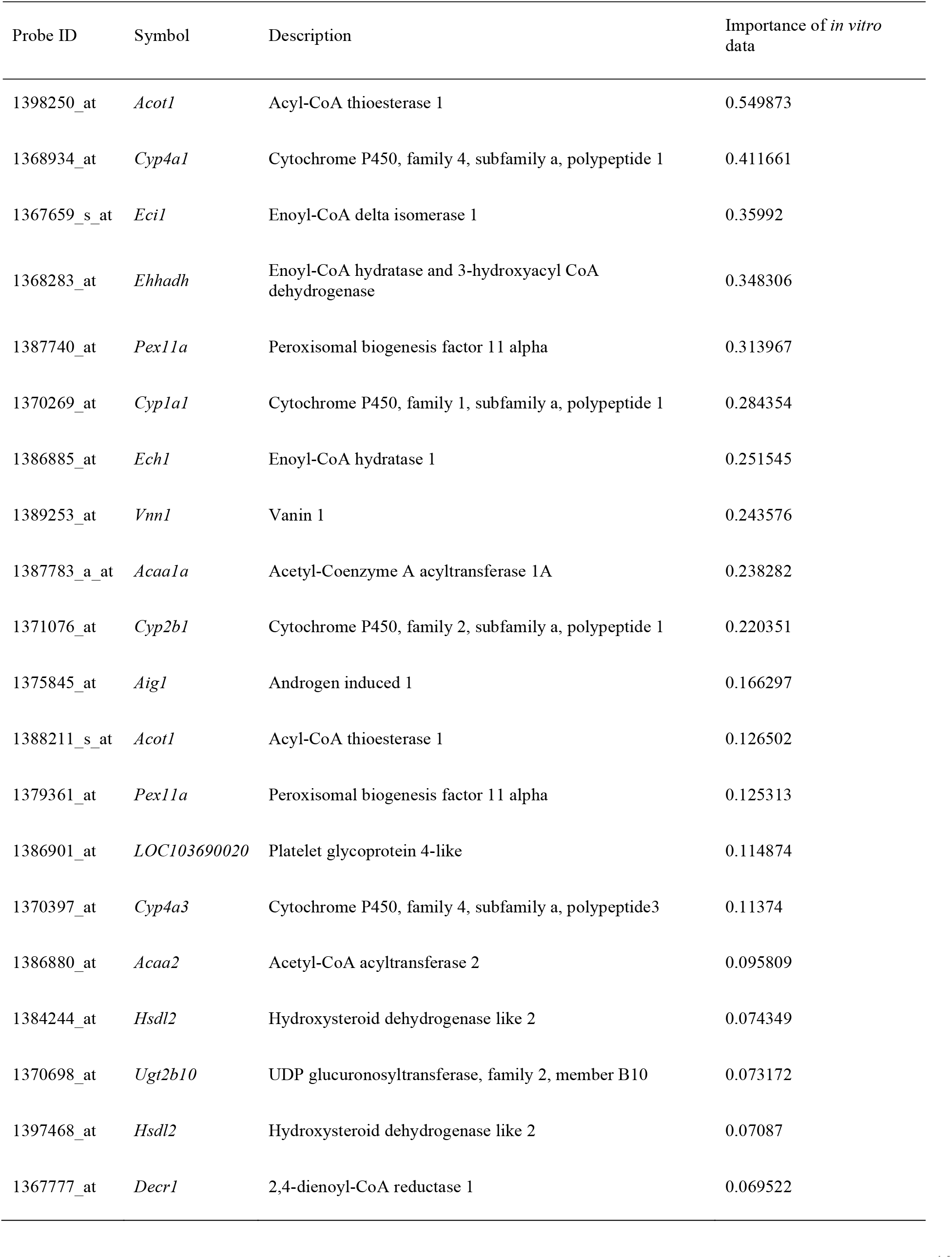
List of top 20 genes with high importance *in vitro* data in the predictive models in RAID.

### 3.3 Quantitative IVIVE performance in the RAID system

To evaluate the RAID performance in terms of gene expression value, RMSEs were calculated for all genes and the genes for which in vitro data had importance in predictive models. Considering RAID would be used in read-across, we compared the RMSEs of RAID data to that of *in vitro* data, which was conventional non-animal test approaches (Figure 6). As a result, RMSEs decreased in RAID, indicating a better performance than what could be obtained using *in vitro* data.

**Figure 6.**
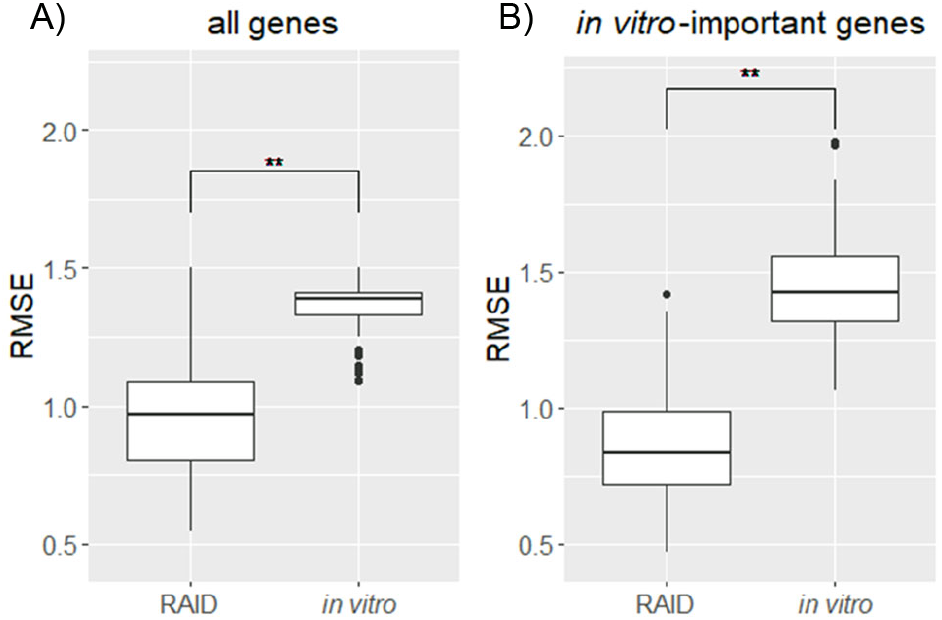
Distribution of RMSEs of a virtual microarray (RAID) and *in vitro* data of A) all genes and B) *in vitro* genes having importance (contribution) in predictive models. **p < 0.01 (Welch’s *t*-test).

### 3.4 Validation of prediction models using external data

In PCA using *in vivo* and RAID data as well as the pathway analysis of PC related genes (Figure 2 and 3), genes related to peroxisome proliferative activity and xenobiotic metabolism activity possibly leading to liver cancer, which were respectively characterized by *Cyp4* induction via PPARa and *Cyp1a* induction via AHR, were identified as key features. Thus, potential *Cyp4a-* and *Cyp1a-*inducers were explored using the knowledge-based approach using the IPA software. Moreover, using the top 30 genes identified using PCA (described in **2.3** section), upstream regulator analysis focusing on chemical substances was performed and 20 chemicals were identified. Finally, a total of 21 chemicals (potential *Cyp1a* inducers: 10 chemicals, potential *Cyp4a* inducers: 11 chemicals) were selected as candidates for external validation and subjected to RAID analyses (Table 4). Substances already present in the TG-GATE (training sets) or had uncalculated chemical descriptors data were excluded.

**Table 4.** List of chemical substances used for external validation of RAID system.

| Name | CAS No. | Name in PCA plot |
| --- | --- | --- |
| *Potential Cyp1a inducers* |  |  |
| 2,3,4,7,8-pentachlorodibenzofuran | 57117-31-4 | Pentachlorodibenzofuran |
| 3,4,5,3',4'-pentachlorobiphenyl | 57465-28-8 | Pentachlorobiphenyl |
| 3-methylcholanthrene | 56-49-5 | Methylcholanthrene |
| 9,10-dimethyl-1,2-benzanthracene | 57-97-6 | Dimethylbenzanthracene |
| Benzo(a)pyrene | 50-32-8 | Benzo(a)pyrene |
| Dexamethasone | 8054-59-9 | Dexamethasone |
| Genistein | 446-72-0 | Genistein |
| 2,2',4,4'-tetrachlorobiphenyl | 1336-36-3 | Tetrachlorobiphenyl |
| Quercetin | 117-39-5 | Quercetin |
| Resveratrol | 501-36-0 | Resveratrol |
| Thiabendazole | 148-79-8 | Thiabendazole |
| *Potential Cyp4a inducers* |  |  |
| Streptozotocin | 18883-66-4 | Streptozotocin |
| 2-ethylhexanol | 104-76-7 | Ethylhexanol |
| Di(2-ethylhexyl) phthalate | 117-81-7 | Di(2-ethylhexyl)_phthalate |
| Clofenapate | 21340-68-1 | Clofenapate |
| Clofibric acid | 882-09-7 | Clofibric_acid |
| Ciprofibrate | 52214-84-3 | Ciprofibrate |
| Nafenopin | 3771-19-5 | Nafenopin |
| TO-901317 | 293754-55-9 | TO-901317 |
| Acetaminophen | 719293-04-6 | Acetaminophen |
| Diltiazem | 33286-22-5 | Diltiazem |

For the PCA analysis, approximately half of the substances were plotted with positive PC scores, which is in consistence with the direction expected from the training data set for both potential *Cyp1a-* and *Cyp4a-*inducers (Figure 7). Lastly, pentachlorobiphenyl, polychlorinated biphenyls, and pentachlorodibenzofuran were isolated as *Cyp1a-*inducers, whereas nafenopin, ciprofibrate, and di(2-ethylhexyl) phthalate were isolated as *Cyp4a-*inducers.

**Figure 7.**
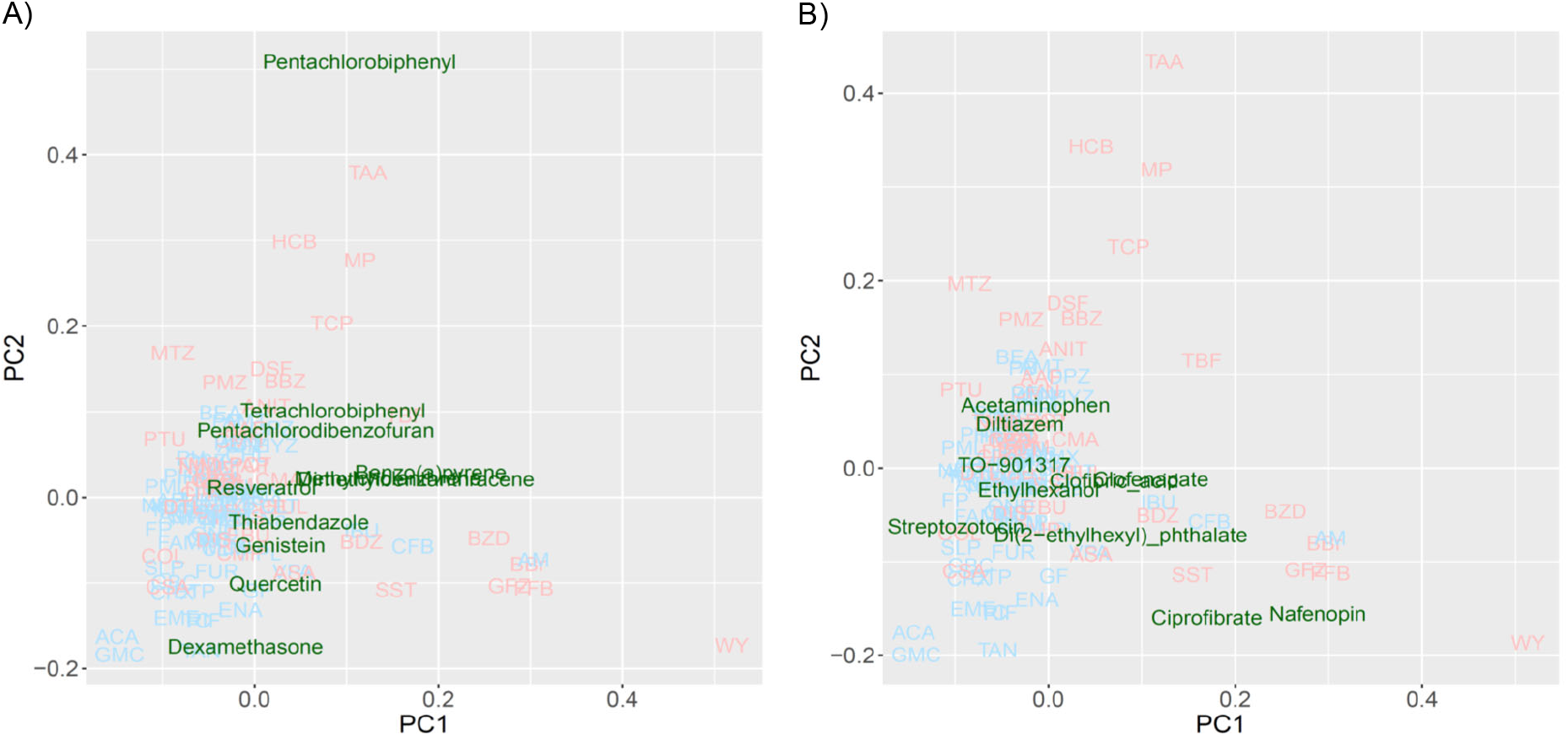
Read-across using PCA plot of external data predicted by a virtual microarray (RAID). A) *Cyp1a* and B) *Cyp4a* inducing chemical substances were analyzed for validation.

## 4 Discussions

The transcriptome data signatures derived from RAID (the virtual microarray) system were in good agreement with those of *in vivo* data, and the technology provided an understanding of the features of hepatotoxic substances based on the toxicological mechanism interpretation. The mechanism of action of the two characteristic toxic substances separated using PCA analysis was shown to be achieved through *Cyp4a* induction via PPARa and *Cyp1a* induction via AHR (pathway and gene ontology analysis). The PPARa-induced drug toxicity requires species differentiation considerations (Ito et al., 2006) and AHR-induced drugs raise safety concerns during developmental periods (Qin et al., 2019). Therefore, predicting the involvement of these nuclear receptors and induction of metabolic enzymes is critical for understanding the molecular initiating events and the key events associated with adverse outcome pathway. RAID enables the prediction of gene expression levels; thus, exhibiting properties required for next generation risk assessment methods.

The 1^st^ substance group (TAA, MP, and HCB), representing toxic substances commonly differentiated from non-toxic substances using PCA on *in vivo* and RAID data, has been reported to have carcinogenicity with metabolic activation (Uehara et al., 2008; Hajovsky et al., 2012; US. HSS., 2015). Furthermore, they have been shown to activate xenobiotic related receptors, such as AHR inducing *Cyp1a* (Ushel et al., 2002; Yamashita et al., 2014; Clara et al., 2015). Moreover, *in vivo* transcriptome data in this study showed that TAA, MP, and HCP induce *Cyp1a* activation. AHR is known for mediating the toxicity and tumor promoting properties despite the mechanism through which AHR activates carcinogenesis remains to be elucidated (Safe et al., 2013; Murray et al., 2014).

The 2^nd^ substance group (WY, FFB, BBr, and GFZ) includes fibrates which are recognized as PPARa agonists (Schoonjans et al., 1996), implying that induction of *Cyp4a* via PPARa and perturbation of lipid-related genes are involved as a series of key events. Although another fibrate included in training data, clofibrate (CFB), was classified as a non-toxic substance according to no serum chemistry findings from a previous study, CFB was shown to act as a PPARa agonist inducing peroxisomal proliferation on hepatocyte (Low et al., 2011) and was plotted around the 2^nd^ group in PCA. Sustained activation of PPARa signaling and induction of enzymes, such as CYP4A, to increased fatty acid oxidation contributes to sustained oxidative stress in liver. These changes lead to liver cell damage as hypertrophy and proliferation which contribute to the development of hepatocellular carcinomas (Parimal et al., 2013).

From the perspective of capturing individual gene responses, RAID was able to detect gene expressions related to major drug metabolism responses in *in vivo* more broadly (more common principal component related gene number; Figure 4) and quantitatively (less RMSE value; Figure 6) than *in vitro*. The 36 genes that were commonly related to principal components of *in vivo* and RAID data contained genes that were known to be involved in drug metabolism and hepatotoxicity. In addition to the genes described above (*Cyp1a* and *Cyp4a*), *Acot1* acts as an auxiliary enzyme in the oxidation process of various lipids in peroxisomes (Hunt et al., 2012). Furthermore, *Vnn1* is expressed by the centrilobular hepatocytes and is involved in lipid and xenobiotic metabolism (Bartucci et al., 2019), whereas *Pex11a (Peroxisomal biogenesis factor 11 alpha)* is involved in peroxisome maintenance and proliferation associated with dyslipidemia (Chen et al., 2018). All of these genes are known as PPARa target genes (Rakhshandehroo et al., 2010; Lake et al., 2016). Thus, these features indicate that RAID can predict possible toxicity by taking into account a broader range of mechanisms than the range of *in vitro* data. Indeed, the *in vivo* changes detected using the *in vitro* data were limited (Figure 4), and the PCA showed most of the differentially expressed genes were associated with irrelevant non-physiological conditions. Thus, the IVIVE effect combining QSAR technique and *in vitro* data would allow for more precise predictions through denoising this type of *in vitro* specific biological responses.

*In vitro* data contribute to accurate gene expression predictions that could not be achieved with QSAR alone (Figure 2D). *In vitro* data contributed to the prediction of the mechanism shown in Figure 5. The biological mechanisms related to metabolic processes were consistent with the key mechanisms of characteristic hepatotoxic substances described above, which indicates that *in vitro* data contributes to the precise predictions obtained using RAID. In addition, whether *in vitro* responses were observed in the suggested mode of action predicted by the RAID system or not is an important point in term of weight of evidence. This study provides valuable evidence supporting that transcriptome data should be considered in light of previous reports indicating that *in vitro* data does not necessarily reflect *in vivo* conditions (Tamura et al., 2006; Sutherland et al., 2016). Simultaneously, *in vitro* studies focusing on a specific mechanism should consider the external validity of their findings and whether the findings reflect *in vivo* situations.

Evaluating the read-across performance using external substances, such as 3,4,5,3’,4’-pentachlorobiphenyl, 2,2’,4,4’-tetrachlorobiphenyl (a type of polychlorinated biphenyl) and pentachlorodibenzofuran (dioxin-like compounds) (Figure 7A), which are known as IARC group 1 carcinogens and *Cyp1a1* inducers (EPA, 1996; Walker et al., 2005; National Toxicology Program, 2006), were separated as toxic-substances. Additionally, benzo(a)pyrene, 3-methylcholanthrene, and 9,10-dimethyl-1,2-benzanthracene plotted apart from origin of coordinates (PC1 = 0 and PC2 = 0) and are polycyclic aromatic hydrocarbons inducing *Cyp1a1* (Moorthy et al., 2007; Pushparajah et al., 2008). Non-carcinogenic chemical substances, such as foods components or preservatives, were positioned near the origin, second quadrant or third quadrant, indicating low risk. Furthermore, substances interacting with *Cyp4a* (Figure 7B), such as ciprofibrate, nafenopine, clofenapate, clofibric acid, and di(2-ethylhexyl) phthalate, which are plotted in the area of the 2^nd^ substance group (PC1 > 0), are also known as PPARa agonist (Bocos et al., 1995; Roberts et al., 2002; Yadetie et al., 2003; Currie et al., 2005; Pyper et al., 2010). Chemicals that were not characterized by the PC1 component (PC1 < 0) are not hyperlipidemia drugs. These results suggest that the RAID system effectively classifies substances that based on the mode of action as well as the strength of toxicity, and ultimately contributes to precise read-across. Thus, the RAID system provides a new method for read-across in line with IATA that should be called “a virtual functional read-across”. Here, we showed that compounds without high structural similarities might have similar toxicological properties, and our new approach interpreted the shared mechanism of action. This means that RAID considers the qualitative and quantitative similarities of biological responses, which was one of the major issues of QSAR-based read-across. The structural similarities of TAA, MP, and HCB observe using correlation coefficient of the chemical descriptor used for the predictive model and the maximum common substructure (MCS) similarities with the Tanimoto coefficient is less than 0.5; however, the homology of RAID and *in vivo* data is as high as 0.8. Furthermore, achieving such an accurate read-across without using *in vitro* data will provide a new perspective on the structural information-based predictions.

PCA analysis was used to understand the features of substances to predict the modes of action and identify biologically similar compounds for read-across in this study. Hence, focusing on certain specific toxicity, discriminant analysis, classifier model, or biomarker analysis might improve the separation of toxic substances. Indeed, the use of RAID data instead of experimental transcriptome data would achieve previously reported biomarker-based classification without using animals. For example, Liu et al. (2017) indicated that certain genes associated with hepatocellular hypertrophy and hepato-carcinogenesis, and markers, such as *Cyp1a1*, *Acot1*, *Stac3 (SH3 and cysteine rich domain 3)*, and *Hdc (Histidine decarboxylase),* which were correctly evaluated in the present study to characterize hepatotoxic compounds in PCA. Similarly, the constructed RAID system could be applied to previous studies to predict carcinogenicity or estimate transcriptional benchmark dose by toxicogenomics analysis of short term *in vivo* studies (Ellinger-ziegelbauer et al., 2008; Thomas et al., 2013; Matsumoto et al., 2014; Kawamoto et al., 2017).

One important issue that should be considered in toxicological evaluation using the RAID system is consideration of species differences. The RAID system provides mechanistic insights on repeated-dose toxicity in animal models; however, since some species differences have observed, the suggested mode of action and the corresponding molecules need to be confirmed by toxicologists. Moreover, evaluation on RAID usefulness for various toxicities is required.

The present approach integrates QSAR and IVIVE and will contribute to other areas of research, such as drug repositioning, which recently attracted attentions towards pharmaceuticals that are available on the market and might be repurposed for new diseases (Jourdan et al., 2020). However, the previously proposed methodologies (Iwata et al., 2018; Lippmann et al., 2018; Zhu et al., 2020; He et al., 2021) have a room for improving the IVIVE aspect of *in vivo* prediction. Thus, our system provides an alternative to screen candidate drugs and explore new biologically similar drugs at a low cost.

In conclusion, we developed a virtual DNA microarray system that quantitatively predicts *in vivo* gene expression profiles based on the chemical structure and/or *in vitro* transcriptome data. Estimated transcriptomes are considered scientifically relevant from PCA data interpretation as well as pathway and GO analysis. Based on its external validation, our system works as an alternative test for repeated dose toxicity tests with toxicogenomic analysis enabling IVIVE and mechanism estimation. Although our technology might have limited applicability domain due to the small data size of chemical substances and their characteristic (using hepatotoxic substances), the concept of the virtual microarray analysis contributes to 3Rs and might benefit every future animal testing.

## 6 Author Contributions

YA and HH contributed to conception and design of the study. YA and HH constructed in silico models, performed enrichment analyses, interpret the biological meanings of the models, and contributed to statistical analyses. HH collected the datasets from TG-GATE. HH and MY supervised this project. YA and HH drafted the manuscript. All authors contributed to manuscript writing, confirmed the final version of the manuscript, and agreed to the contents.

## 8 Acknowledgments

We thank Dr. Osamu Morita, Dr. Kaede Miyata, and Mr. Yasuaki Inoue for their helpful suggestions and valuable discussions to the present study.

## 10 Data Availability Statement

Publicly available datasets were analyzed in this study. This data can be found here: https://toxico.nibiohn.go.jp/open-tggates/english/search.html.

